# Feeding status modulates diel vertical migration of zooplankton via effects on circadian rhythms

**DOI:** 10.1101/2024.07.12.603318

**Authors:** Cory A. Berger, Ann M. Tarrant

## Abstract

Diel vertical migration (DVM) of aquatic animals is arguably the largest migration on Earth, and occurs each day in most marine and freshwater ecosystems. DVM is influenced directly by factors such as light, food, and predator abundance, but is also regulated by internal circadian clocks. Untangling the mechanistic controls of DVM, and the relative importance of external versus endogenous cues, has been hampered by a lack of experimental systems. Here, we leverage advances in animal tracking software to develop an imaging system allowing us to quantify the positions of dozens of individual copepods (*Acartia tonsa*) at sub-second resolution over multiple days. We use this approach to characterize group-level diel behavioral patterns at much higher resolution than has previously been possible. We find that behavioral rhythms entrain to light cycles and regular feeding and persist during constant conditions (darkness, no feeding), indicating true circadian regulation and establishing *A. tonsa* as an experimental system for DVM. We test the hypothesis that food availability impacts DVM by acting as an entraining cue for circadian rhythms. Daytime-restricted feeding weakens circadian behavioral rhythms compared to nighttime-restricted feeding, illustrating that food availability can impact DVM indirectly via effects on internal clocks in zooplankton. Our results provide a detailed view of zooplankton populations over diel cycles, establish a mechanism by which temporal variation in food supply can affect zooplankton migrations, and lay the groundwork for new experimental studies of circadian behavior.

## 1 Introduction

Diel vertical migration (DVM) of aquatic animals is arguably the largest migration on Earth by biomass (Brierley, 2014) and occurs every day in marine and freshwater systems across the globe. At ecosystem scales, DVM contributes to the spatiotemporal structure of food webs and drives species interactions (Hays, 2003; Kelly et al., 2019). At a global scale, DVM influences the carbon budget and biogeochemical cycling due to the active transport of carbon and nutrients to depth (Archibald et al., 2019; Brierley, 2014; Kelly et al., 2019; Steinberg et al., 2000). These large migrations are caused by the movements of individual animals whose behavior is influenced by external cues and endogenous (circadian) mechanisms. We do not yet understand the relative contributions of external vs. endogenous control of DVM behavior, nor how the influence of these factors on individual behavior scales up to affect group-level migrations.

The most common pattern of DVM is the nighttime ascent of herbivorous zooplankton, which is ultimately driven by adaptive trade-offs between grazing activity and predation risk (Tarling et al., 2003). Herbivorous zooplankton eat phytoplankton that primarily occur near the surface of the water column, but are vulnerable to visual predators in the daylit surface waters. Therefore, many grazers take refuge in deeper and darker waters during the day, only ascending to feed at night. “Reverse DVM” can occur when the migrators’ predators are not visual and themselves perform nighttime DVM (e.g., chaetognaths; Tarling et al., 2003).

This adaptive framework for DVM is well-supported empirically, but does not directly explain why the timing, magnitude, and patterns of migration differ among populations and across time and space. Nor does it explain the extensive inter-individual variability that is often present in DVM behavior (i.e. one population often contains both migrators and non-migrators; Hays et al., 2001). Environmental factors such as food availability, predator abundance, and light, as well as internal factors like circadian clocks, all influence the migratory behavior of zooplankton in natural populations, but experiments are required to disentangle the relative contributions of these factors to behavior patterns.

Light is certainly the best-studied proximal driver of DVM. Changes in relative light levels are a commonly-observed trigger for DVM, consistent with bouts of migration around dawn and dusk (Brierley, 2014). It is clear that light levels strongly influence DVM in general (Cohen & Forward Jr., 2009), but the specific mechanism(s) by which zooplankton sense and respond to light are controversial. The main hypotheses are that 1) migrators track a preferred isolume, 2) they respond to changes in light intensity, or 3) they sense and respond to the relative rate of change of light, irrespective of absolute irradiance. These hypotheses are difficult to test because it is challenging to draw mechanistic conclusions from population-level observations. For instance, vertical zooplankton distributions are, in some cases, closely associated with isolumes, which has been suggested as strong evidence for the isolume hypothesis (Cohen & Forward Jr., 2009; Hobbs et al., 2021; Sainmont et al., 2014). However, this community-level pattern does not imply that individual animals follow preferred light levels. In fact, all of these mechanistic hypotheses (isolume, rate of change, and relative rate of change) could in principle result in similar nocturnal DVM patterns (Richards et al., 1996)—and none of them can fully explain all attributes of DVM.

Mesocosm and laboratory experiments show that food availability modulates the amount of time that migrating animals spend at the surface, with starved animals generally spending more time at the surface (Calaban & Makarewicz, 1982; Clarke, 1932; Forward & Hettler, 1992; Johnsen & Jakobsen, 1987). This is interpretable as increased foraging effort and tolerance of predation risk. Field studies have also observed more time at the surface during low-food conditions (Huntley & Brooks, 1982) and more time at depth during high-food conditions (Dagg et al., 1997)—although the results of this latter study appear to be driven by the presence of phytoplankton at depth rather than satiation of grazers. The situation is complicated by the fact that food availability can influence behavior in multiple ways: first, by affecting hunger/satiety; second, by directly attracting grazers to the food source; and third, by affecting circadian clocks. In any case, nutritional status is a likely driver of variability in DVM among individuals of the same population, since it influences the costs and benefits of migrating (Hays et al., 2001).

Zooplankton also modulate their migration patterns in direct response to predators (or predator cues). Unsurprisingly, zooplankton are more likely to migrate when predator cues are abundant in the water column (e.g. Bollens et al., 1992; Forward & Hettler, 1992; Frost & Bollens, 1992), and some studies have observed a lack of zooplankton DVM in the absence of predators (Bollens & Frost, 1989; Loose, 1993; Ohman, 1990; Pangle & Peacor, 2006). Mechanistically, these results are driven by behavioral plasticity of zooplankton in response to predators, rather than differential mortality imposed by predators on behaviorally-fixed non-migrators (Bollens & Frost, 1991). It appears that predator detection primarily occurs via chemical cues, although visual or mechanical detection may also play a role (Bollens et al., 1994).

Some patterns of DVM cannot be explained by simple effects of light or other environmental factors. Most strikingly, DVM can occur during the polar night (Berge et al., 2009) and at depths below the photic zone (van Haren & Compton, 2013), when diel light cues are weak or absent. Other, more subtle, migration patterns are also somewhat mysterious, including the pattern of halting ascent before reaching the surface (Richards et al., 1996) and “midnight sinking” followed by a second nighttime ascent (Cohen & Forward, 2002). Although these behaviors could result from unobserved or complex responses to external cues, endogenous rhythmicity provides an attractive explanation for aspects of DVM that occur without obvious external signals. Circadian regulation of zooplankton DVM is also supported indirectly by observations of clock gene expression and other physiological rhythms in migrating animals (e.g. Häfker et al., 2017; Maas et al., 2024; Tarrant et al., 2021).

The strongest evidence for the role of circadian clocks in DVM comes from laboratory experiments that demonstrate endogenous swimming rhythms in various crustaceans (Cohen & Forward, 2005; Duchêne & Queiroga, 2001; Gaten et al., 2008; Gerhardt et al., 2006; Häfker et al., 2017; López-Duarte & Tankersley, 2007; Piccolin et al., 2020; Zeng & Naylor, 1996). These studies have observed DVM-like rhythms in the vertical distributions of zooplankton that occur even in constant conditions (i.e. free-running), thus demonstrating circadian regulation of vertical swimming behavior. However, these studies share two main limitations: 1) they have generally used field-collected animals with unknown environmental histories (as most zooplankton are difficult to culture); and 2) they have used low-resolution behavioral metrics involving manual counting of animals by human observers at discrete intervals. To gain mechanistic insight into the relative contributions of environmental factors and endogenous rhythms to DVM, we require experimental systems that use cultured species and automated behavioral measurements with high spatiotemporal resolution.

To address these limitations, we develop the cosmopolitan copepod *Acartia tonsa* as a laboratory model for zooplankton DVM. Recent advances in multi-animal tracking software based on machine learning enable quantification of the 2-dimensional positions of dozens of individual copepods over multiple days using off-the-shelf camera equipment and open-source software. We use this methodology to 1) show that *Acartia* exhibits DVM-like rhythms in the lab, and 2) test the hypothesis that the timing of food availability impacts DVM by modulating circadian rhythms. Consistent with studies in other copepods, we identify light-synchronized DVM-like rhythms on small (<35 cm) spatial scales that persist in constant conditions. Our findings show that the relative timing of environmental signals can facilitate or inhibit group-level migratory behavior. In general, these tracking data provide a high-resolution view of copepod movement ecology, shedding light on how population-level patterns emerge from the behaviors of individual animals.

## 2 Materials and methods

### 2.1 Animal culture

*Acartia tonsa* were derived from a laboratory population originally collected from Long Island Sound in Groton, CT, USA. Copepods were maintained in full-strength, 0.35 µm-filtered seawater at 20 °C in a 14:10 light-dark (LD) cycle and fed a monoalgal diet of *Rhodomonas marina*. Algae were grown at 20 °C in a 14:10 LD cycle, with f/2 medium (without silica) in full-strength, 0.35 µm-filtered seawater that was sterilized by microwaving (Keller et al., 1988). The main population of *Acartia* was maintained in 2-3 mixed cultures containing all life stages, in volumes of approximately 3 L. Full water changes were performed every two weeks. Periodically, eggs were harvested and hatched separately, allowing us to rear animals that were approximately synchronized in development.

Approximate food concentrations were determined by counting cells with a hemocytometer and converting from cells to carbon concentration based on an estimated carbon content for *R. marina* of 34 pg C per cell (Mullin et al., 1966; Olenina et al., 2006). The target food concentration fed to copepods each day was ∼100 µg L^−1^ C. Once the culture was reproducing successfully and the population was stable, we stopped counting algae cells each day and instead selected algae volumes based on appearance (color) of the culture; thus, food concentrations should be viewed as approximate.

### 2.2 Experimental setup

The imaging setup consisted of two thin, vertical plexiglass tanks with internal dimensions of 40 × 11× 4 cm (1.76 L; Figs. 1A, 1B). Tanks were filled with filtered seawater to a height of 35 cm and lit from above by visible strip lights (full-spectrum white light, 6500 kelvin LEDs; intensity was roughly 250 lux at the top of tanks) on a programmed LD cycle and continuously by infrared (IR) LEDs (850 nm). Tanks were imaged with Basler daA3840-45um cameras (8.3 megapixel resolution) with IR-pass filters and recorded at 2 frames per second (fps). To record experiments, 20-100 animals retained on a 350 µm filter (which includes adults and larger copepodite stages; see Discussion) were transferred to the tanks and two trials were recorded simultaneously.

**Figure 1:**
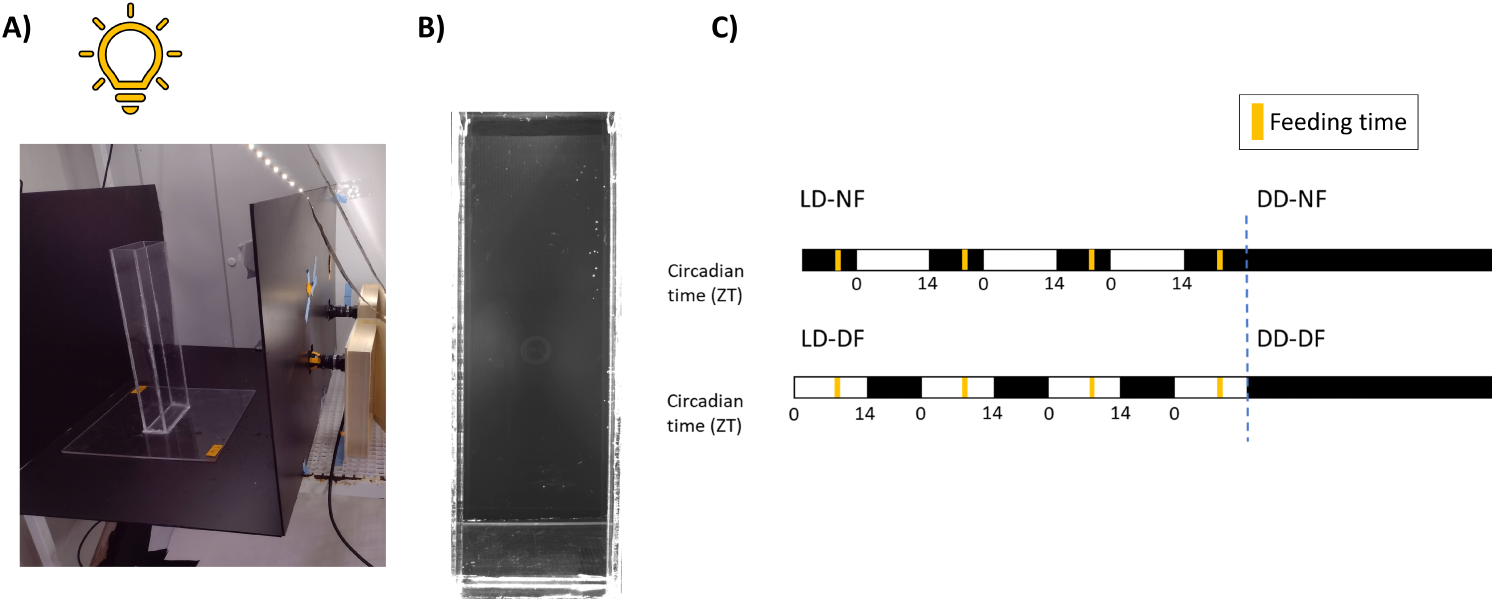
Imaging setup and experimental design. A) Photograph of imaging setup, consisting of a plexiglass tank lit from above and imaged from the side with an IR camera. B) Example frame of camera footage. Note that animals are not visible. C) Schematic of experimental design. To perform time-restricted feeding experiments, animals were reared in one of two 14:10 LD cycles, which were offset from one another by 12 hours. The animals were then fed at the same clock time (yellow bars), achieving nighttime feeding in LD-NF and daytime feeding in LD-DF. ZT0 always refers to lights-on. Thus, ZT0 in LD-NF occurs at the same time as ZT12 in LD-DF, and vice versa. For free-running trials, animals from LD-NF and LD-DF conditions were transferred to constant darkness with no feeding (DD-NF, dashed blue line).

In total, we recorded behavior trials under four conditions, which are shown in Fig. 1C. We recorded trials during a light-dark cycle with nighttime feeding (LD-NF) or with daytime feeding (LD-DF); animals were fed during these recordings. We also recorded trials with animals that were reared in LD-NF or LD-DF but transferred to constant darkness with no feeding (DD-NF and DD-DF). We define lights-on as ZT0 (“zeitgeber time”), and lights-off occurred at ZT14. In order to create the nighttime feeding (NF) and daytime feeding (DF) groups, animals were maintained in inverted LD cycles and fed at the same clock time, corresponding to either ZT20 (NF) or ZT8 (DF). Copepods for feeding experiments were derived from the same initial cohort of eggs, which were collected from the main population, split, and reared from hatching in either NF or DF. For each feeding trial, one NF group and one DF group were recorded simultaneously in the two tanks, thus allowing for direct comparisons between trials. The first of these trials were recorded two weeks after hatching, and subsequent trials were recorded over the next several weeks using the same initial populations.

We wanted to minimize handling time and damage to *Acartia*, which are small and fragile. Therefore, we did not precisely count the number of animals placed into each trial, and there is variability in the density of individuals within each experiment. To back-estimate the number of individuals in each trial, we calculated the maximum number of individuals tracked in at least 0.01% of frames (∼50 frames in a 72 h video). This metric was chosen instead of the maximum number of individuals tracked in a single frame to avoid biases due to inaccurate tracking in a small number of frames.

### 2.3 Tracking software and processing

Animal tracking was performed using TRex v1.1.6 (Walter & Couzin, 2021), a machine learning program designed for efficient tracking of many individuals at a time. We used the following parameters for video processing (blob size range=[0.001,0.009], correct luminance=true, use closing=true, use adaptive threshold=true, threshold=14) and tracking (blob size ranges=[0.0007,0.007], categories min sample images=10, gpu max sample gb=0.5, huge timestamp seconds=2, recognition enable=false, track max individuals=250, track max reassign time=4, track max speed=2, track speed decay=0, track threshold=21, track trusted probability = 0.25). These parameters were selected after manual troubleshooting to obtain reliable tracking of copepods.

The main source of error in this analysis occurs at the tracking step, because copepods can be lost or obscured (false negatives), especially along the edges of the tank, or mistakenly confounded with bubbles or reflections (false positives). False positives are more serious than false negatives, and we presume that not all individuals are visible in any given frame. We employed two approaches to identify high-confidence copepod trajectories. First, we used the machine learning categorization functionality of TRex to categorize tracked points as either copepods or noise. Within the software interface, TRex extracts video clips of tracked objects, which the user can then manually label. On a dataset of 9787 manually-labelled frames, TRex achieved 94% training accuracy in categorizing frames as copepod or noise. When applied to the full dataset, TRex categorized frames as copepod, noise, or undetermined, reported per frame and as an average for each trajectory. For subsequent analyses, we used frames that were either 1) categorized as copepods, or 2) categorized as undetermined within a trajectory whose average category was copepod. Second, we applied post-hoc filters to define active copepods. Since *Acartia* cannot remain still in the water column (they either swim or passively sink), we required a minimum movement speed of 0.005 cm s^−1^, averaged over 5 s sliding windows; we also split trajectories into new tracks if speeds exceeded 0.5 cm per frame. In order to improve trajectory continuity, we also interpolated single frames that were dropped within otherwise continuous trajectories. We only analyzed trajectories that fulfilled the movement criteria for at least 10 consecutive frames.

To account for slight differences in camera and tank positioning between trials, we measured the distance in pixels between sides of the tank (a distance of 11 cm), and used these normalization factors to convert distances and speeds from pixels to mm. For each trajectory, we estimated the size of the individual based on its average size (in mm^2^) across all frames. These size estimates cannot be considered reliable measurements of body size because 1) they do not account for the ±10% variation due to movement in the Z dimension, and 2) the bounding boxes used by the software do not accurately capture the true outline of the copepod. Bounding boxes could be overestimated if the software draws the box around the long antennae, or (more likely) underestimated if the software only picks up part of the animal. Nonetheless, these values should capture genuine differences in the size distribution between trials, assuming that there are no systematic biases in these estimates between videos. We assigned individuals to four size classes based on the quartiles of the overall size distribution.

### 2.4 Behavior metrics

In each frame, we obtained position and velocity estimates of tracked individuals from the above trajectories. For velocity estimates, we smoothed individual trajectories with a Savitzky-Golay filter of order 7 (’TrajSmoothSG’ function of the ’trajr’ R package). We then calculated various group-level behavior metrics in each frame (Table 1).

**Table 1:** List of group behavior metrics.

| **Metric** | **Definition** |
| --- | --- |
| Median depth | Median of Y coordinate values |
| Mean depth/Centroid Y | Mean of Y coordinate values |
| Centroid X | Mean of X coordinate values |
| Depth standard deviation | Standard deviation of Y coordinate values |
| Depth inter-quartile range (IQR) | Range between the 0.25 and 0.75 quantiles of Y coordinate values |
| Centroid speed | Speed of the centroid (mean X and Y coordinates) in $\text{cm s}^{-1}$, smoothed with a Savitzky-Golay filter of order 11 |
| Mean speed | Mean individual speed, in $\text{cm s}^{-1}$ |
| Speed inter-quartile range (IQR) | Range between the 0.25 and 0.75 quantiles of individual speeds |
| Polarization | Absolute value of mean individual headings (Tunstrøm et al., 2013). Ranges from 0 (totally unaligned) to 1 (all headings are identical) |
| Group spacing | Mean pairwise distance between all individuals |
| Nearest-neighbour distance | Mean distance between pairs of nearest-neighbours |

In addition to the metrics in Table 1, we calculated curl in order to measure the rotation of the velocity vector field. To calculate this vector field, we used the “oce” and “sf” R packages to divide the tank into a 10×10 grid, calculate mean X- and Y-velocity components in each cell, interpolate missing cells using the Barnes algorithm (’interpBarnes’), and calculate curl for each cell and on average across the entire vector field. This was performed within hourly bins in order to capture longer-term circulation dynamics.

### 2.5 Statistical analyses

To quantify rhythmicity, we used Lomb-Scargle periodograms (LSP) within the ’lomb’ R package, with significance assessed by randomization and a p-value cutoff of 0.01. To test for rhythmicity within each trial, we averaged a given behavior metric in 10-minute bins and used the ’randlsp’ function with ofac=50 to scan for periods between 20 h – 28 h. We excluded the first 6 h of each trial from rhythmicity analysis in order to minimize artifacts from acclimation to the tank. To test for average rhythmicity across an experimental condition (LD-NF, DD-NF, LD-DF, or DD-DF), we averaged data into hourly bins, scaled and centered the hourly data within each trial, averaged bins across trials, and tested for rhythmicity with ’randlsp’ as above. We used periodogram power as a measure of the strength of rhythmicity.

We performed cross-correlation analysis between behavior metrics using the ’prewhiten’ and ’ccf’ functions of the ’TSA’ and ’tseries’ R packages. First, we averaged each metric into hourly bins in each time series. For each time series and pair of metrics, we “prewhitened” the time series by removing autocorrelation from one metric and applying the same decorrelating filter to the other (Dean & Dunsmuir, 2016) using ’prewhiten’, which selects an autoregressive model based on AIC minimization. We calculated cross-correlations up to a 12 h lag within each trial, and averaged coefficients across all trials. We assessed significance of cross-correlations with 99% confidence intervals derived from the standard error at each lag.

We used linear models to test for differences in non-time series data (e.g. number of individuals per trial) among groups or for associations between variables, using the ’lm’ R function and ’emmeans’ R package. In the case of comparing behavior metrics across size classes, we used the ’lme’ function with random intercepts for each trial.

## 3 Results

### 3.1 Automated behavior tracking of Acartia recapitulates group movement dynamics

Using a laboratory population of *Acartia tonsa*, we developed an assay to track many individual copepods in an automated fashion. For each experimental replicate, we placed ∼20-100 animals into tall, thin tanks (4 × 11 × 40 cm) which were filled with seawater to a height of 35 cm, illuminated from above with IR light (850 nm) and imaged at 2 frames per second (Fig. 1). By minimizing the Z-dimension, we sought to approximate the 2D position of animals in the water column. Using the machine learning software TRex (Walter & Couzin, 2021), we obtained high-confidence individual trajectories with a mean length of ∼12 s across all trials (18 trials total, each with a length of roughly 72 h). Trajectory lengths were limited by constraints such as camera resolution and the loss or obfuscation of animals by bubbles and tank edges, and we chose to prioritize high-confidence tracks rather than longer but lower-confidence trajectories. Therefore, the data consist of a series of overlapping but relatively brief trajectories such that, in any given frame, we obtain XY coordinates and velocities of a subset of individuals. The mean number of tracked individuals per frame across all trials was 6.99. Although we could not track individual animals for extended periods of time, these data allow us to quantify group-level parameters at sub-second resolution over multiple days.

We first showed that our data processing pipeline captures group-level behaviors that are visible with the naked eye. Visual inspection of videos identified situations where *Acartia* moved in a “vortex” pattern—i.e., they generally ascended on one side of the tank and descended on the other. This pattern often persisted for multiple days. This type of group swimming behavior has been observed in other zooplankton in small-volume cultures, and is probably the result of collision-avoidance behavior in constrained spaces (Mach & Schweitzer, 2007; see also Discussion). Regardless of the underlying reason, it provides a useful ground truth for us to evaluate the accuracy of our tracking data.

Encouragingly, the vortex pattern was evident in our processed data. We calculated velocity vector fields over the entire tank, representing the average movement of copepods over a given window of time. Fig. 2 provides an example of a trial in which anti-clockwise rotation was visible in the velocity field averaged over a 72 h video. We can quantify this rotation with curl, which represents the strength of rotation in a vector field. In this case, the average curl was positive throughout the video (Fig. 2D). In other trials, curl was consistently negative, indicating clockwise rotation, or roughly zero, indicating no mean rotation (Table S2). We provide a more detailed analysis of this vortex behavior in a subsequent section, including its variability over diel timescales. For now, we note that this finding demonstrates the ability of our pipeline to accurately describe group-level behaviors in *Acartia*.

**Figure 2:**
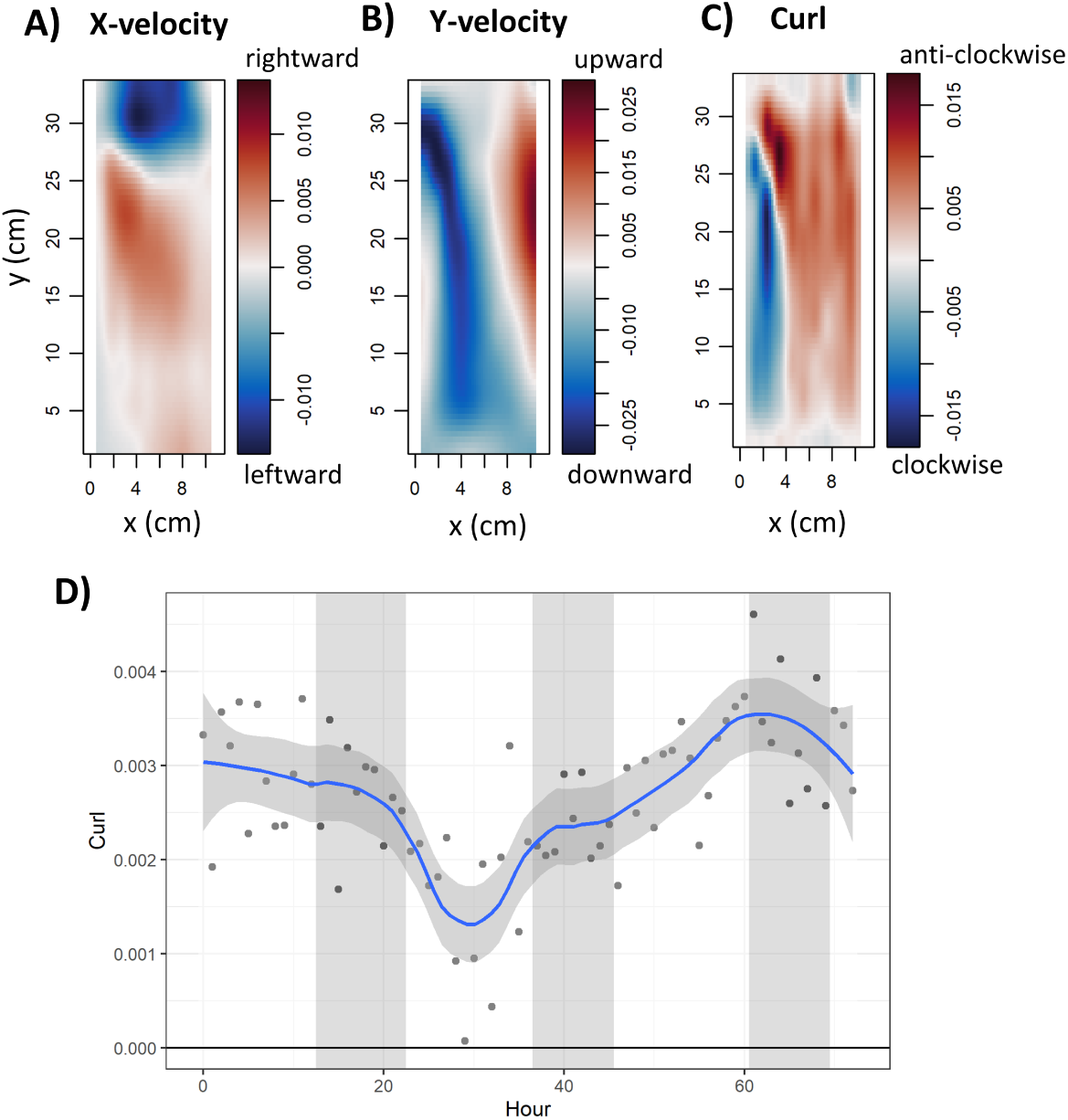
Example of *Acartia* performing vortex-like swimming behavior in an enclosed container. Trial 0621-B; see Table 2 for conditions of each trial). A) X velocities averaged over 72 h. Positive values (red) indicate rightward movement. B) Y velocities averaged over 72 h. Positive values (red) indicate upward movement. Anticlockwise rotation is visible in panels A and B. C) Curl averaged over 72 h. Positive values (red) indicate anti-clockwise rotation. D) Curl calculated within hourly bins and averaged over the entire tank. In this case, curl is consistently positive, indicating anti-clockwise rotation. White/grey rectangles indicate lights-on/lights-off, the blue line is a loess smooth of the data, and the shaded area is the 95% confidence interval.

### 3.2 Acartia perform DVM-like behavior in laboratory conditions

Animals were initially maintained in a 14:10 LD cycle and fed during subjective night (ZT20, with lights-on defined as ZT0). We recorded six 72 h trials in these conditions (LD-NF), and 5 out of 6 trials displayed significant diel rhythmicity in mean and median population depth (LSP, randomization test, p<0.01, period between 20 h – 28 h). See Table 2 for list of trials and accompanying statistics. The overall pattern was a peak in height just before subjective dusk, followed by a lower distribution for most of the night and early day (Fig. 3A). There was some variation in the timings of these rhythms between trials: one trial had a peak later in the night (trial 0621-A), and in others, the ascent occurred earlier in the day (trials 0621-B and 0814-B). Overall, however, this behavior appears to recapitulate the nighttime ascent aspect of classic DVM, indicating a tendency to ascend in the water column prior to dusk. It is worth noting that *Acartia* are positively phototactic, yet the observed diel depth changes cannot be explained by either positive or negative phototaxis. This shows that *Acartia* repeatably display diel changes in depth on small spatial scales in laboratory conditions.

**Table 2:** Trial statistics. Numbers in trial name indicate recording date (e.g., 0621-A and 0621-B were recorded simultaneously on June 21, 2023). Phase was calculated by fitting a linear model with a sinusoidal fit and a period of 24 h (’lm’ function) to binned hourly median depth data. n/a indicates trials with no significant rhythm (LSP, p>0.01). Trials from 0709 or later were produced from the same initial clutch of eggs hatched on June 23, 2023.

| Trial | Condition | LSP<br>power,<br>median<br>depth | Rhythmic<br>depth<br>phase<br>(h) | Rhythmic<br>depth<br>period<br>(h) | Estimated<br>individ-<br>uals | Median<br>size<br>(mm<sup>2</sup>) |
| --- | --- | --- | --- | --- | --- | --- |
| 0621-A | LD-NF | 0.123 | 17 | 20.7 | 27 | 0.239 |
| 0621-B | LD-NF | 0.113 | 7 | 24.6 | 24 | 0.267 |
| 0709-A | LD-NF | 0.105 | n/a | n/a | 68 | 0.268 |
| 0719-A | LD-NF | 0.117 | 10 | 22.6 | 32 | 0.262 |
| 0726-B | LD-NF | 0.106 | 12 | 23.5 | 30 | 0.277 |
| 0814-B | LD-NF | 0.315 | 9 | 22.4 | 33 | 0.263 |
| 0630-A | DD-NF | 0.054 | n/a | n/a | 18 | 0.211 |
| 0630-B | DD-NF | 0.057 | 18 | 21 | 13 | 0.278 |
| 0706-A | DD-NF | 0.234 | 13 | 22.5 | 20 | 0.261 |
| 0706-B | DD-NF | 0.248 | 18 | 23.1 | 52 | 0.280 |
| 0802-A | DD-NF | 0.063 | n/a | n/a | 77 | 0.234 |
| 0808-B | DD-NF | 0.335 | 8 | 23.5 | 17 | 0.227 |
| 0719-B | LD-DF | 0.016 | n/a | n/a | 27 | 0.256 |
| 0726-A | LD-DF | 0.033 | n/a | n/a | 60 | 0.264 |
| 0814-A | LD-DF | 0.027 | n/a | n/a | 42 | 0.269 |
| 0802-B | DD-DF | 0.179 | n/a | 28.1 | 22 | 0.248 |
| 0808-A | DD-DF | 0.295 | n/a | 30.4 | 29 | 0.218 |

**Figure 3:**
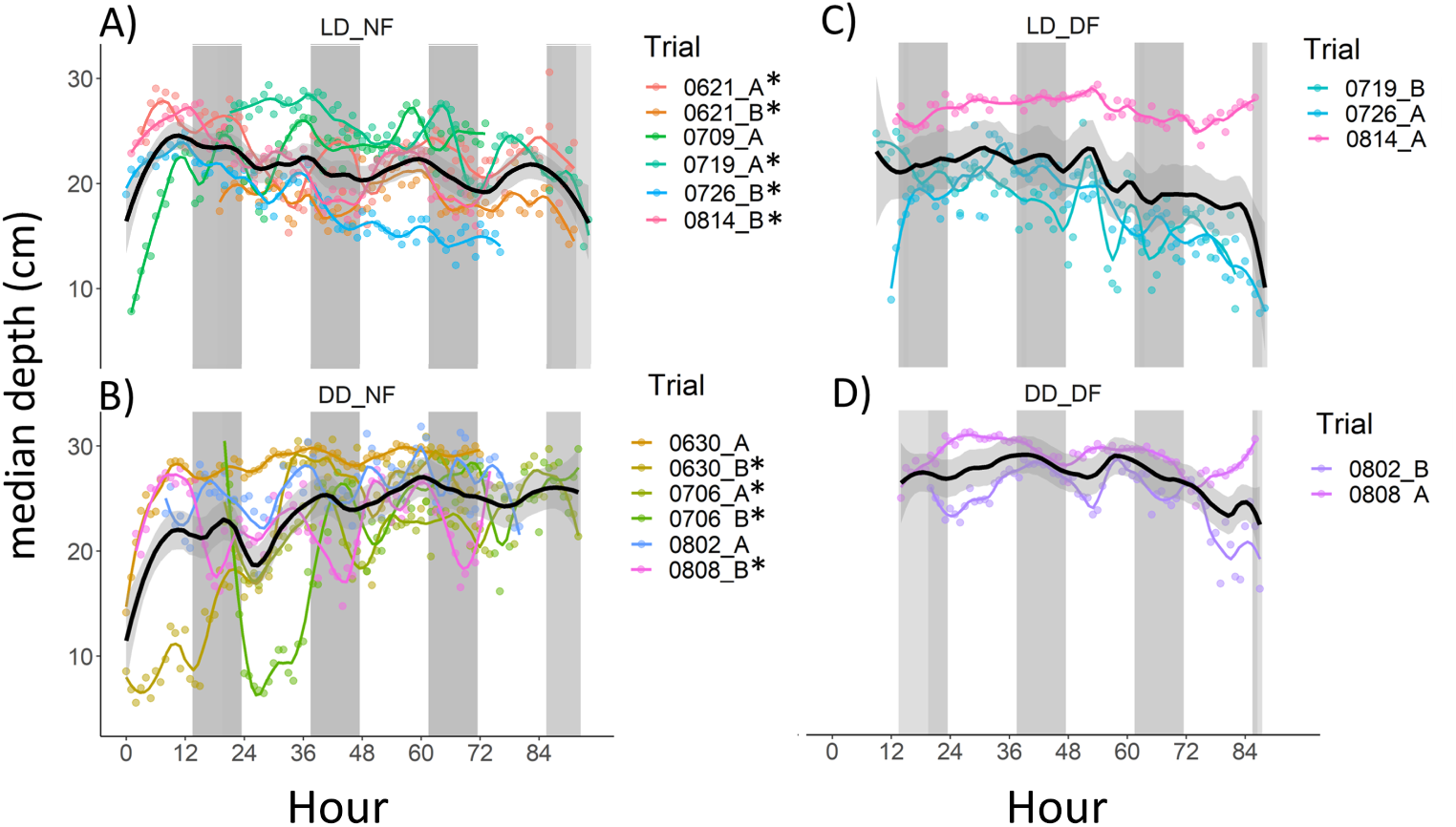
Median depth over time. Points represent hourly averages and colored lines represent loess smooths for each trial. Black lines represent loess smooths of all data within each condition, and shaded areas represent 95% confidence intervals. A) LD-NF. B) DD-NF. C) LD-DF. D) DD-DF. Asterisks indicate trials with significant rhythmicity between 20 h – 28 h (LSP, p<0.01). White/grey rectangles represent lights-on/lights-off (A,B) or subjective day/night (C,D).

Diel rhythms in depth persisted during constant conditions of darkness and no feeding (DD-NF; Fig. 3B), showing that DVM-like behavior is under circadian control and not simply driven directly by light or food availability. During constant conditions, 4 out of 6 trials had diel rhythms in mean and median depth (Table 2). The overall pattern of ascent prior to subjective dusk was still visible and statistically significant when averaged across all trials (LSP, p<0.01). There was no significant difference in the strength of rhythmicity (LSP power) between LD-NF and DD-NF trials (lm, p=0.8).

### 3.3 Feeding experiments

We next conducted time-restricted feeding experiments in which copepods were reared in 14:10 LD cycles and fed either at ZT20 (night-time feeding, LD-NF) or at ZT8 (day-time feeding, LD-DF); see Fig. 1 for a schematic of the experimental design. For these experiments, we used paired cohorts derived from the same initial group of eggs. Eggs were collected from the main population and divided into two groups, each reared on a different feeding/light regime; all other conditions were identical. Thus, we can specifically contrast pairs of trials sampled from the same genetic background and at the same time point, relative to their initial hatching. One trial (0709-B) was discarded because it was inadvertently exposed to incorrect light cues.

None of the 3 LD-DF trials had detectable rhythms in mean or median depth (Fig. 3C). In contrast, all 3 paired LD-NF trials had significant rhythmicity, with an ascent during late day (Fig. 4A). This indicates that diel behavioral rhythms are driven not only by the light cycle, but also depend on the time of food availability.

**Figure 4:**
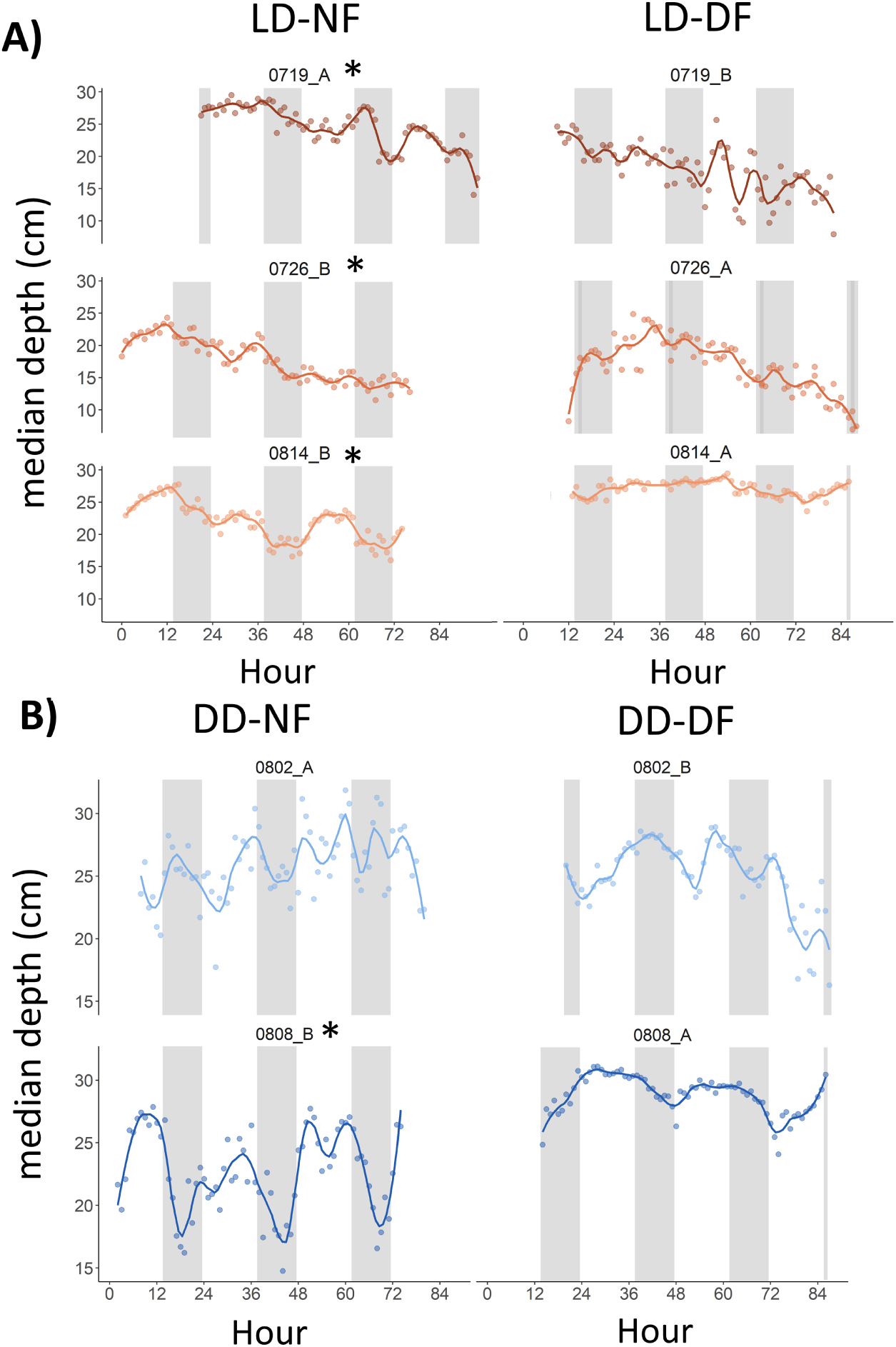
Daytime-restricted feeding alters DVM-like swimming behavior in *Acartia*. An egg cohort was divided into either nighttime feeding (NF, left column) or daytime feeding (DF) groups. Each row represents a pair of trials recorded simultaneously. A) Median depth during LD-NF (left) and LD-DF (right). B) Median depth during DDNF (left) and DD-DF (right). White/grey rectangles indicate lights-on/lights-off (A) or subjective day/night (B). Asterisks indicate22trials with significant diel rhythmicity (LSP, p<0.01).

We also recorded trials for NF and DF groups during free-running (constant darkness, no feeding). Both of the DD-DF trials had depth rhythms with periods longer than 28 h—just beyond our criteria for “circadian” (Fig. 3D). In contrast, no other rhythmic time series had a period close to 28 h (Table 2). One of the paired DD-NF trials (trial 0808-B) had clear circadian rhythmicity (Fig. 4B). The other paired trial (trial 0802-A) was not rhythmic, but most animals were apparently dead/untrackable by the second day (see Discussion).

### 3.4 Other behavior metrics

Having demonstrated circadian rhythms in depth distribution, we next examined how other behavioral metrics for *Acartia* varied over diel timescales (Table S1). After normalizing and averaging across trials, LD-NF groups exhibited diel rhythmicity in centroid X position, centroid speed, mean speed, speed inter-quartile range (IQR), group spacing, nearest-neighbour distance, and curl (Fig. S1). DD-NF exhibited rhythmicity in centroid X, centroid speed, mean speed (marginal, p=0.01), and nearest-neighbour distance. LD-DF exhibited rhythmicity in speed IQR and polarization, while DD-DF was not rhythmic in any of these examined metrics.

We performed cross-correlation analysis to further understand relationships between behavior metrics. Across all trials, we observed significant positive cross-correlations (with lag=0) of median depth with mean speed and IQR speed; and negative cross-correlations of median depth with centroid speed, group spacing, nearest-neighbour distance, depth standard deviation, depth IQR, and polarization.

This analysis allows us to form a qualitative description of DVM in *Acartia*. As the population ascended during late day, the mean movement speed of individuals was also somewhat faster, along with increased variability among the speeds of different individuals. This suggests that only a subset of individuals increased their activity. At the same time, the overall spacing of the group became closer/less dispersed, as group spacing and nearest-neighbour distance were at their lowest. The depth distribution was also narrower at this time, indicating compression specifically in the vertical dimension. To further explore the effects of DVM on the depth distribution, we looked at rhythmicity of different quantiles of the depth distribution. Rhythmicity was stronger within lower (deeper) depth quantiles, with peak statistical rhythmicity occurring at roughly the 0.4 quantile for LD-NF (Fig. S2). This indicates that the deeper portion of the population undergoes greater diel variation in depth, and that DVM—in this specific context—is due to shoaling of deeper individuals (Fig. 5). Finally, centroid speed and polarization were low during the ascent, indicating reduced coordination in the headings of different individuals. Again, this suggests that only a subset of animals performed the migratory behaviour, rather than all individuals uniformly.

**Figure 5:**
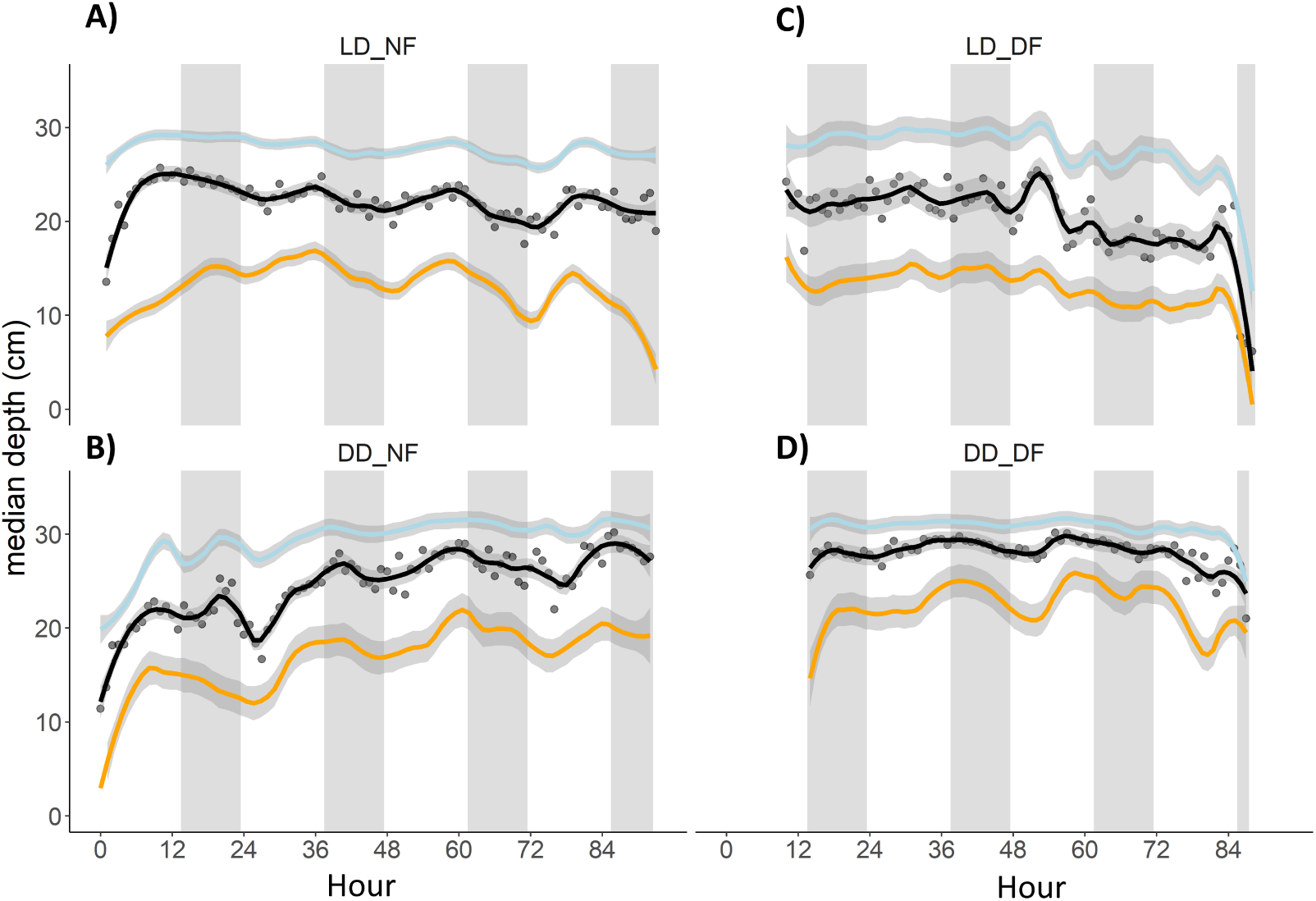
DVM reflects a shoaling of deeper individuals. Points represent hourly averages of median depth across all trials. Lines represent loess smooths of the 0.75 (blue), 0.5/median (black), and 0.25 depth quantiles (orange). Shaded areas represent 95% confidence intervals. A) LD-NF. Note that the 0.25 quantile undergoes greater depth variation than the 0.75 quantile. B) DD-NF. C) LD-DF. D) DD-DF. White/grey rectangles indicate lights-on/lights-off or subjective day/night.

### 3.5 Curl

As discussed previously, many trials exhibited a pattern of “vortex” swimming behavior. We calculated curl within hourly bins to examine the temporal variability of this pattern. Qualitatively, 5 out of 18 trials had consistent positive curl, 3 trials had consistent negative curl, and the remaining trials had curl with no clear direction on average (Table S2, Fig. S3). Although curl was generally consistent within trials, some trials exhibited strong diel rhythmicity in curl, including 5 LD-NF trials, 1 DD-NF trial, and 1 LD-DF trial (Table S2). Three of these trials switched between clockwise and anti-clockwise rotation in a diel manner (see Fig. S4 for an example). Curl was rhythmic on average across LD-NF trials, with stronger rotation at night. Curl was also overall higher in LD-NF compared to other groups, when calculated within hourly bins (lm, p<0.0001). Unlike other behavior metrics, rhythmicity in curl may be light-driven rather than circadian, because 1) it did not occur during free-run, and 2) it seems to directly vary with lights-on/lights-off rather than being anticipatory.

We considered that variation in curl may simply reflect individual activity, as animals that are less active would result in weaker rotational velocities. However, we found no relationship between curl and mean speed across hourly bins, whether considered across the entire dataset or just within rhythmic curl trials. Therefore, this behavior is not simply a product of swimming speeds. Additionally, we found no relationship between the number of individuals per trial and curl (lm with random effects for trial, p=0.5), nor with the strength of rhythmicity (LSP power) of curl.

### 3.6 Heterogeneity among trials

Developmental stage, sex, size, and density may all contribute to heterogeneity in diel behaviour among trials. Although we cannot control for theses factors in a rigorous way, we examined whether differences in density or size structure contributed to variation among trials. The estimated number of individuals per trial did not differ systematically between groups (lm, p=0.7). Additionally, number of individuals in a trial was not related to the strength of rhythmicity (Fig. S5A).

We assigned sizes to individual trajectories based on the average number of pixels occupied by the tracked individual across all frames. The over-all size structure was a smooth, right-tailed distribution with a median of 0.255 mm^2^ (as noted in the Methods, these sizes should only be considered rough estimates rather than precise measurements).

Size structure differed across trials (ANOVA, p<0.05), but not systematically between conditions (ANOVA, p=0.28). The largest difference in median size (considering all individual tracks irrespective of length) was between 0630-A (0.211 mm^2^) and 0706-B (0.280 mm^2^; Table 2). Across all trials, there was an average size increase of 5.4% over 72 h (lm with random intercepts for trials, p<0.01), likely reflecting growth and molting throughout the recordings. There was no relationship between median size per trial and periodogram power (Fig. S5B), indicating that differences in rhythmicity between groups were not due to differences in size structure.

Nonetheless, we examined whether there were behavioral differences between size classes within and across trials. To do this, we assigned each individual to one of four size classes based on the quartiles of the overall size distribution, which we have named from A (largest) to D (smallest). There were some consistent differences among size classes. Unsurprisingly, larger size classes moved faster and had higher IQR speed (lm, p<0.01; Table 3). Additionally, larger animals were consistently found higher in the water column (Fig. 6), and occupied narrower vertical ranges (smaller SD Y and IQR Y). Smaller animals had higher within-class polarization and faster centroid speeds.

**Figure 6:**
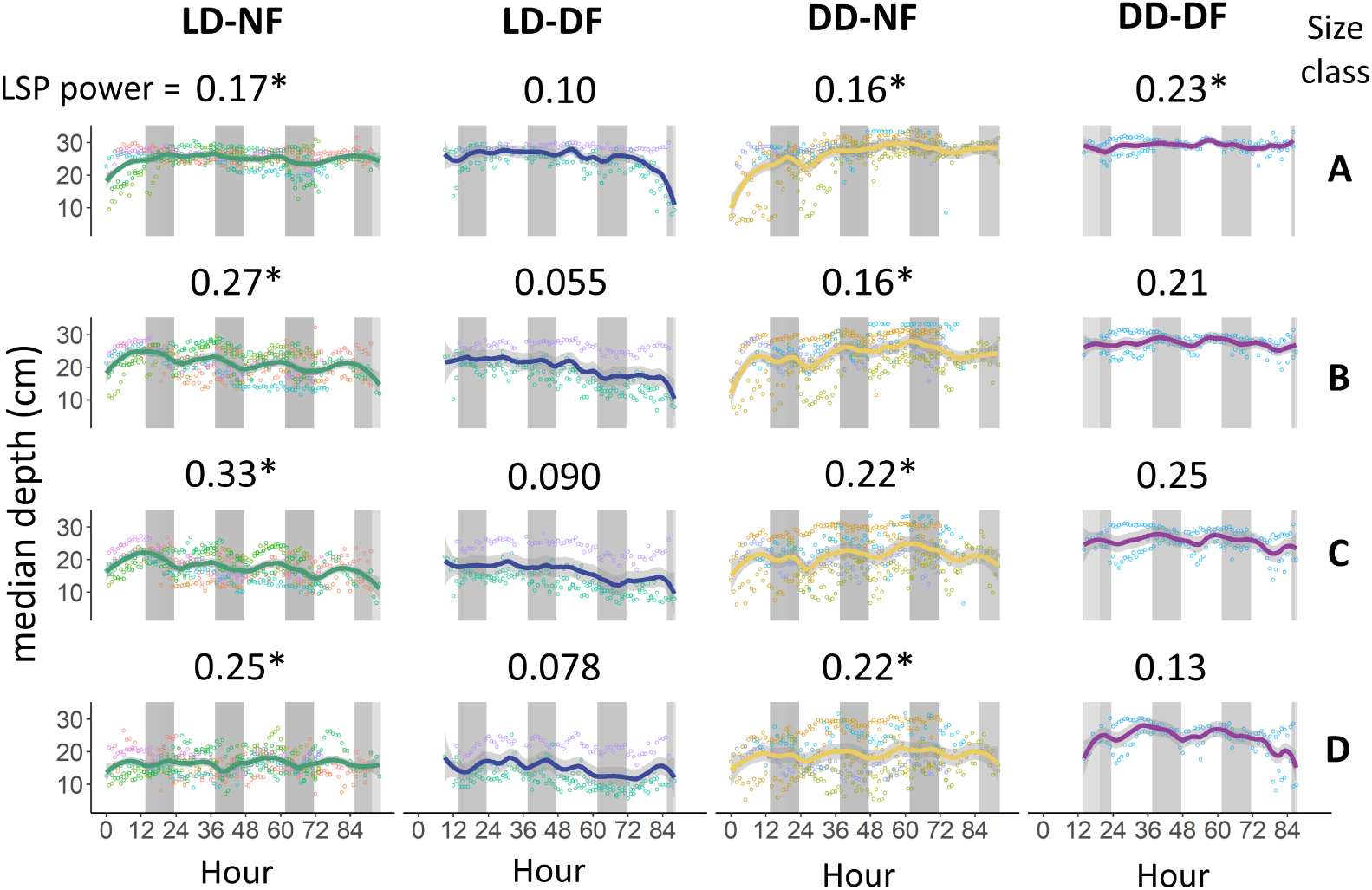
Size classes differ in depth distribution and rhythmicity. Size classes range from the largest (A, top row) to smallest (D, bottom row) 25% bins. Points represent hourly averages of median depth in a trial. Lines represent loess smooths of hourly data averaged across all trials. Shaded areas represent 95% confidence intervals. White/grey rectangles indicate lights-on/lights-off or subjective day/night. Numbers above panels indicate power of mean time series between 20 h – 28 h; asterisks indicate significant rhythmicity between 20 h – 28 h; for DD-DF size classes B-D, periodogram peaks were >28 h.

**Table 3:** Mean behavior metrics within conditions and size classes.

| Condition | Size<br>class* | median<br>depth<br>(cm) | mean<br>speed<br>(cm/s) | IQR<br>speed<br>(cm/s) | IQR<br>depth<br>(cm) | Polarization<br>(within<br>class) | Centroid<br>speed<br>(cm/s) |
| --- | --- | --- | --- | --- | --- | --- | --- |
| LD-NF | A | 24.5 | 0.050 | 0.020 | 3.64 | 0.72 | 0.72 |
| LD-NF | B | 21.4 | 0.043 | 0.018 | 4.58 | 0.71 | 0.91 |
| LD-NF | C | 17.9 | 0.035 | 0.012 | 4.39 | 0.77 | 1.1 |
| LD-NF | D | 16.4 | 0.026 | 0.0080 | 3.91 | 0.82 | 1.4 |
| LD-DF | A | 25.6 | 0.045 | 0.019 | 3.42 | 0.74 | 0.64 |
| LD-DF | B | 20.1 | 0.041 | 0.016 | 4.44 | 0.75 | 0.77 |
| LD-DF | C | 16.4 | 0.033 | 0.010 | 3.95 | 0.80 | 0.90 |
| LD-DF | D | 14.9 | 0.026 | 0.0062 | 3.1 | 0.86 | 1.1 |
| DD-NF | A | 26.3 | 0.030 | 0.0082 | 1.98 | 0.85 | 0.50 |
| DD-NF | B | 24.2 | 0.027 | 0.0076 | 2.81 | 0.84 | 0.62 |
| DD-NF | C | 21.4 | 0.025 | 0.0060 | 2.71 | 0.86 | 0.79 |
| DD-NF | D | 19.2 | 0.022 | 0.0041 | 2.06 | 0.90 | 0.92 |
| DD-DF | A | 29.0 | 0.036 | 0.011 | 0.69 | 0.79 | 0.42 |
| DD-DF | B | 27.4 | 0.034 | 0.015 | 2.91 | 0.69 | 0.52 |
| DD-DF | C | 25.6 | 0.029 | 0.013 | 3.31 | 0.69 | 0.66 |
| DD-DF | D | 24.2 | 0.023 | 0.0096 | 3.78 | 0.72 | 0.93 |

\*Individuals were assigned to one of four size classes corresponding to the quartiles of the overall size distribution, ranging from A (largest 25%) to D (smallest 25%).

The results of our feeding experiments were robust to comparisons within any given size class (Fig. 6). Overall, rhythmicity in depth was strongest in size class C (between the first and second quartiles). Among LD-NF trials, all four size classes had strong and significant rhythmicity in median depth, with size class C having the highest power, followed by B, D, and A. In DD-NF, all four size classes were also rhythmic. In LD-DF, none of the size classes were rhythmic. In DD-DF, size class A had a significant period at 27.5 h, technically within our range of circadian periods; the other size classes all had periods longer than 28 h.

## 4 Discussion

We present a methodology for the high-resolution tracking of groups of zooplankton over multiple days, and use this approach to quantify circadian swimming rhythms in *Acartia tonsa*. Calanoid copepods such as *Acartia* occupy central positions in pelagic and coastal food webs and are dominant vertical migrators by biomass in many ecosystems (Darnis et al., 2017; Paffenhöfer & Stearns, 1988). As with other marine and estuarine copepods, we found that *Acartia* displays endogenous rhythms in depth that reflect the night-time ascent aspect of diel vertical migration (DVM). These rhythms depend not only on a light cycle, but also on time of feeding. Daytime-restricted feeding (DF) appears to abolish diel rhythms in depth and alters underlying circadian rhythmicity. The relative ease of collecting this type of data with long-term cultured populations and modern behavior tracking software will enable further manipulative experiments to clarify the endogenous and environmental drivers of DVM.

For simplicity, we refer to depth rhythms observed over small spatial scales (centimeters) in the laboratory as “DVM”, but acknowledge that this behavior only represents a proxy for natural, large-scale DVM.

### 4.1 What is the role of circadian clocks in DVM?

Across multiple generations and experimental replicates, we observed population ascents prior to subjective dusk corresponding to the expected pattern of DVM nighttime ascent. The fact that this behavior persisted during free-running conditions is evidence that DVM in *Acartia* is under circadian control, as in other copepods (Cohen & Forward, 2005; Häfker et al., 2017). These endogenous rhythms occurred in a long-term laboratory culture, indicating that circadian rhythms in DVM do not require prior entrainment to natural environmental conditions nor the presence of predators. DVM depends on interactions between endogenous rhythms and external signals such as light and food availability, which are difficult to predict in natural environments. Copepods in our study descended throughout the night, rather than remaining close to the surface as would be expected under “classic” DVM. Thus, it is likely that circadian clocks only regulate certain aspects of DVM behavior. Our study and others (Cohen & Forward, 2005; Rudjakov, 1970) indicate that ascent near dusk and descent near dawn are the most likely aspects of DVM to be under circadian control, while other aspects (e.g. residency time at the surface) may depend on direct environmental signals and trade-offs due to nutritional status, food availability, and predator abundance.

### 4.2 Detailed descriptions of group-level diel behavior in Acartia

#### 4.2.1 Variation in migratory behavior among individuals

Individual trajectories provide both position and velocity data, giving us a more detailed quantitative picture of diel zooplankton behavior than has heretofore been possible. Previous laboratory studies of DVM have—out of necessity—reduced population depth to low-resolution observations such as the number of individuals in the top half of the water column. In reality, population depth is a distribution, and DVM may involve changes in grouplevel parameters other than mean/median depth. Previous laboratory studies have not been able to distinguish whether DVM results from uniform movements of the entire population, or from the preferential movement of some individuals. We found that DVM in *Acartia* (in this context) involved more pronounced migrations by deeper individuals, while the upper portion of the population stayed relatively constant over time. This affected the overall group distribution, as *Acartia* became vertically compressed and closer together during the pre-dusk ascent. Thus, migratory behavior of subsets of the population can contribute to diel variation in the overall distribution and density of the group, and probably in the intensity of intra-species interactions. Depth changes were accompanied by increased mean activity and increased variability in activity among individuals, further indicating that DVM was not performed uniformly by all individuals.

We observed depth stratification of *Acartia* based on size, with larger size classes consistently occurring higher in the water column. This was true across all trials and was independent of diel variation in depth. This may correspond to niche partitioning among developmental stages. Ontogenetic niche partitioning is common among zooplankton in both natural and laboratory populations—although it is more common for older stages to occur deeper, rather than shallower, likely reflecting their increased vulnerability to visual predators (Kobari & Ikeda, 2001; Landry, 1978; Leibold et al., 1994; Mauchline, 1998). In our case, without predators, niche partitioning by depth may reflect competition among stages. The upper water column is probably a more favorable environment, as it is closer to the food source and further away from detritus on the bottom. Adult and late copepodite *Acartia spp.* can cannibalize smaller stages (usually nauplii, but also possibly early copepodites; Camus & Zeng, 2009; Lonsdale et al., 1979), and thus smaller stages may avoid larger copepods. All size classes performed DVM, but DVM was statistically strongest in the 0.25 - 0.5 quantile size range. Due to the observed depth stratification by size, this observation is confounded with the observation of deeper individuals performing stronger DVM. An additional complication is that, before they reach adult stage, *Acartia* molt every 1-2 days (Miller et al., 1977). Therefore, we expect the stage and size distribution to change over the course of the trial, and indeed we observed an increase in individual size over time across all trials, corresponding to an average increase of 5.4% over 72 h. Although the inclusion of mixed life stages in our study is in some sense a limitation, migrating natural populations obviously consist of many life stages. Our observation of rhythmicity across size classes suggests that most copepodite stages of *Acartia* possess endogenous rhythms in DVM.

There may also be behavioral differences between sexes (male *Acartia* are smaller than females and less tolerant of starvation; Finiguerra et al., 2013), but we are not aware of any study of variability in depth distribution or DVM between sexes in *Acartia*.

#### 4.2.2 Vortex swimming behavior

In many trials, we observed a vortex-like swimming pattern involving collective vertical rotation in a single direction. This pattern has been observed in *Daphnia* in the horizontal, rather than vertical, plane (Ordemann et al., 2003), and likely helps minimize collisions while maintaining constant motion (Mach & Schweitzer, 2007; Vollmer et al., 2006). We would therefore expect the direction of rotation to be essentially random, and indeed we observed examples of both clockwise and anti-clockwise rotation. Oddly, we also observed several trials in which there was no clear rotation, and some in which rotation alternated between clockwise to anti-clockwise in a diel fashion.

The factors that contribute to variation in vortex swimming are unknown. It seems that light plays a role, because rotation was significantly stronger during lights-off compared to lights-on in LD-NF trials, apparently causing diel rhythmicity in the strength of rotation. However, there were also freerunning trials with consistent rotation. We found no relationship between the strength of rotation and the speed of individuals, suggesting that factors other than overall activity contribute to the strength of rotation. We also found no relationship between density of *Acartia* and the average strength of rotation in a trial, which is surprising if rotation is in fact driven by collision avoidance.

Although they are not the main focus of our study, these observations illustrate how high-resolution tracking data provide new insights into (and raise new questions about) zooplankton collective behavior. In *Daphnia*, vortex behavior was observed over timescales of minutes (Ordemann et al., 2003), whereas we found that rotation persisted for multiple days. The relevance of this behavior to natural populations is unclear (but see Brown & Gibbons, 2022; Lobel & Randall, 1986).

### 4.3 Implications for our understanding of DVM in natural populations

DVM-like behavior in our study is clearly far removed from the adaptive and environmental framework of natural DVM. Nonetheless, controlled experiments allow us to isolate the effects of specific variables that may affect migration patterns in the field. We provide the first experimental evidence that food availability is a relevant zeitgeber for zooplankton circadian clocks, and that this impacts endogenous swimming rhythms.

Nighttime feeding (NF) is the usual situation for many herbivorous zooplankton, including *Acartia*. We found that daytime feeding (DF) abolished DVM-like behavior. This is similar to other studies of “sensory conflict,” whereby mismatches between two zeitgebers disrupt or weaken normal behavioral rhythms (Berger & Tarrant, 2023; Harper et al., 2016). DF copepods also displayed altered free-running rhythms, with notably longer periods than NF animals (28 h – 30 h). Free-running circadian periods are naturally variable among individuals and species, but typically close to 24 h (23 h – 25 h in *Drosophila*, for example; Srivastava et al., 2019). Exceptionally long or short free-running rhythms suggest weaker entrainment to environmental variables. Thus, our results indicate that food availability impacts diel behavior in zooplankton specifically via entrainment of circadian clocks.

The finding that DF alters circadian swimming rhythms mirrors observations in terrestrial mammals, in which time-restricted feeding can alter behavioral rhythms (Reinke & Asher, 2019; Zhai et al., 2022). DF is commonly used for zooplankton cultures for practical reasons. Our results show that this choice has behavioral consequences, and scientists should consider nighttime feeding if they want to approximate natural conditions. However, NF is far from ubiquitous in zooplankton populations. Various species of copepods can perform reverse DVM as a result of predation by invertebrate predators (Ohman et al., 1983), including nauplii and early copepodites of *Acartia* (Cuker & Watson, 2002; Holliland et al., 2012). In such cases, feeding will primarily occur during the day. Our study provides evidence that this altered feeding regime can affect circadian rhythms and in turn influence diel swimming patterns.

Distinguishing between classic and reverse DVM raises a subtle point. Environmental factors influence behavior, but behavior also controls the environment that an organism experiences. For instance, zooplankton that migrate vertically throughout the day experience a vastly different light regime than if they remained at constant depth. Thus, migratory behavior can feed back on itself: DVM controls the light regime that an individual experiences, which in turn influences DVM behavior. Similarly, animals that feed at a certain time of day due to predator avoidance will have altered endogenous rhythms, which in turn affect their migration patterns. The ultimate outcomes of these interactions are complex, as any change to the circadian clock could not only affect locomotor behavior directly, but also by modulating sensitivity to light or other variables. In general, oceanographers should consider the role of diel signals other than light in contributing to patterns of DVM. These factors may include food and predator abundance, temperature, and oxygen levels.

### 4.4 Limitations and future directions

Various factors may contribute to variability among trials in our study. These include differences in the stage/size structure and density of animals within trials. Since neither of these metrics differed significantly between groups or had a relationship with strength of rhythmicity, our results were not sensitive to these factors. Nonetheless, future studies could test for differences in diel behavior between life stages or in relation to density.

Additionally, we did not directly measure algae concentrations or seawater chemistry, and there may have been variation in these factors across trials or small-scale fluctuations within trials. The repeatability of our results across trials and generations makes this unlikely to affect our conclusions. Still, it would be beneficial to precisely control for nutritional status and test the effects of hunger on swimming rhythms. *Acartia* only live 2-4 days without food, and we expect some mortality during 72 h free-running trials. There was an overall decrease in mean speed over time in free-running trials (lm, p<0.001), but not in LD trials (lm, p=0.35), consistent with reduced activity during starvation. In one case (trial 0802-A), nearly all animals appear to have died by the second day of free-running. However, other free-running trials still had many active individuals on the third day.

Further developments to the tracking methodology would be beneficial. More sophisticated imaging setups could record at higher resolution and in three dimensions, improving tracking accuracy and enabling the use of larger tank dimensions (as behavior may differ on larger spatial scales or with less restrictive dimensions). In principle, improved tracking resolution could distinguish individual identities over multiple days, revealing individual-level variation in migratory behavior.

## Acknowledgements

We are particularly indebted to Drs. Hans Dam and Gihong Park, who generously shared their copepod and phytoplankton cultures, and to Dr. Matt Johnson, who also provided phytoplankton cultures and advice. We thank Phil Alatalo for culturing advice and providing equipment; Dr. Sam Laney for advice about the imaging setup; and Casey Machado and the Dunkworks staff for helping with construction of the experimental tanks.

## Data availability

Raw tracking data, processed trajectory data, and code are freely available on Dryad: https://doi.org/10.5061/dryad.vx0k6dk14.

