## Supplemental File for "Feeding status modulates diel vertical migration of zooplankton via effects on circadian rhythms"

**Table S1**: Rhythmicity of all metrics across experimental groups

| Metric | LD-NF rhythmicity (LSP p-value) | LD-DF rhythmicity | DD-NF rhythmicity | DD-DF rhythmicity |
|---|---|---|---|---|
| Median depth | **<0.001** | 0.097 | **<0.001** | na |
| Mean depth | **<0.001** | 0.082 | **<0.001** | na |
| Centroid X | **<0.001** | 0.067 | **<0.001** | 0.26 |
| Depth SD | 0.014 | 0.36 | 0.19 | 0.019 |
| Depth IQR | 0.027 | 0.288 | 0.46 | 0.05 |
| Centroid speed | **0.004** | 0.504 | **0.008** | na |
| Mean speed | **0.006** | na | 0.01 | 0.63 |
| Speed IQR | **<0.001** | **<0.001** | 0.402 | 0.56 |
| Polarization | **0.007** | **<0.001** | 0.77 | 0.65 |
| Group spacing | **0.001** | 0.15 | 0.076 | na |
| Nearest-neighbour distance | **0.002** | 0.389 | **<0.001** | 0.11 |
| Curl (absolute value) | **0.001** | 0.022 | 0.018 | 0.43 |

Definitions of behaviour metrics are given in Table 1. Bolded cells indicate p-values < 0.01. “na” indicates that the peak periodogram value was outside the interval of 20-28 h. SD: standard deviation; IQR: inter-quartile range.

**Table S2**: Curl information per trial

| Trial | Condition | Curl direction* | Rhythmic curl? | Average curl value** |
|---|---|---|---|---|
| 0621-A | LD-NF | positive | Yes | $8.5 \times 10^{-6}$ |
| 0621-B | LD-NF | positive | No | $2 \times 10^{-3}$ |
| 0709-A | LD-NF | negative | Yes | $-7.9 \times 10^{-4}$ |
| 0719-A | LD-NF | negative | No | $-1.0 \times 10^{-3}$ |
| 0726-B | LD-NF | switches | Yes | $7.7 \times 10^{-5}$ |
| 0814-B | LD-NF | switches | Yes | $3.6 \times 10^{-4}$ |
| 0630-A | DD-NF | negative | No | $-1.0 \times 10^{-3}$ |
| 0630-B | DD-NF | switches | Yes | $-1.8 \times 10^{-4}$ |

| | | | | |
|---|---|---|---|---|
| 0706-A | DD-NF | no clear direction | No | $-1.6 \times 10^{-4}$ |
| 0706-B | DD-NF | no clear direction | No | $8.4 \times 10^{-5}$ |
| 0802-A | DD-NF | no clear direction | No | $-3.5 \times 10^{-4}$ |
| 0808-B | DD-NF | positive | No | $7.4 \times 10^{-4}$ |
| 0719-B | LD-DF | positive | Yes | $2.9 \times 10^{-4}$ |
| 0726-A | LD-DF | no clear direction | No | $3.5 \times 10^{-5}$ |
| 0814-A | LD-DF | positive | No | $1.4 \times 10^{-5}$ |
| 0802-B | DD-DF | positive | Yes | $8.5 \times 10^{-5}$ |
| 0808-A | DD-DF | positive | No | $2 \times 10^{-5}$ |

*Curl direction was assigned qualitatively after manual inspection of plots of hourly curl. "Positive" and "negative" refer to trials with consistent positive or negative curl throughout the trial, "no clear direction" indicates trials with curl that often crosses zero and has no rhythmicity, and "switches" refers to trials with strong rhythmicity, where curl switches between positive and negative with a 24h period.

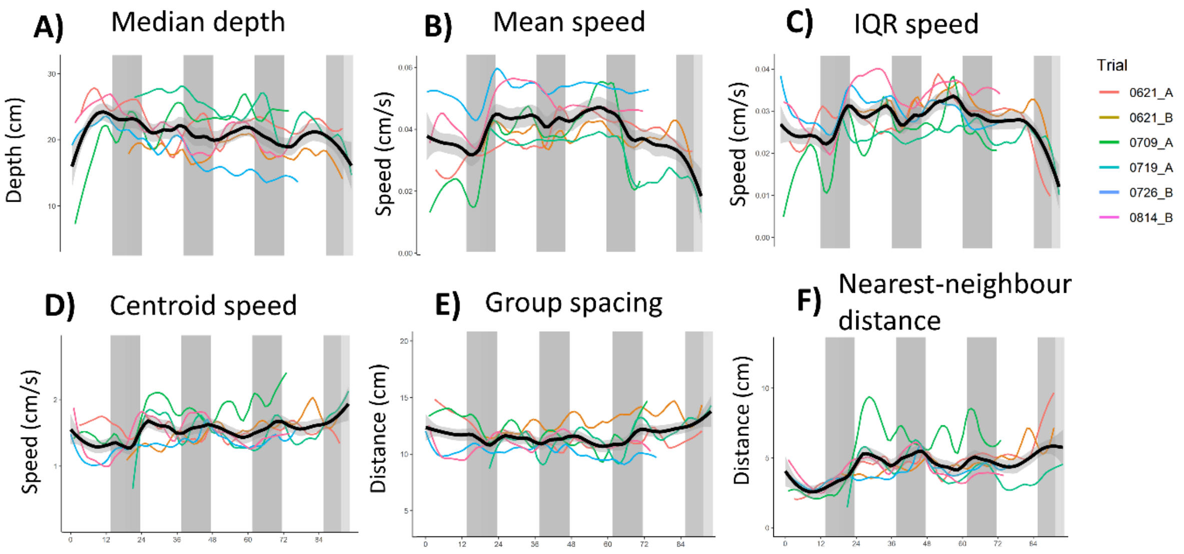


**Figure S1:** Additional behavior metrics with rhythmicity in LD-NF. Colored lines represent loess smooths of hourly averages for each trial (points not shown for clarity). Black lines represent loess smooths of all data within each condition, and shaded areas represent 95% confidence intervals. Only LD-NF trials are shown. A) Median depth. B) Mean speed. C) Speed inter-quartile range (IQR). D) Centroid speed. E) Group spacing (mean pairwise spacing). F) Nearest-neighbor distance (mean distance to nearest neighbor). White/grey rectangles represent lights-on/lights-off.

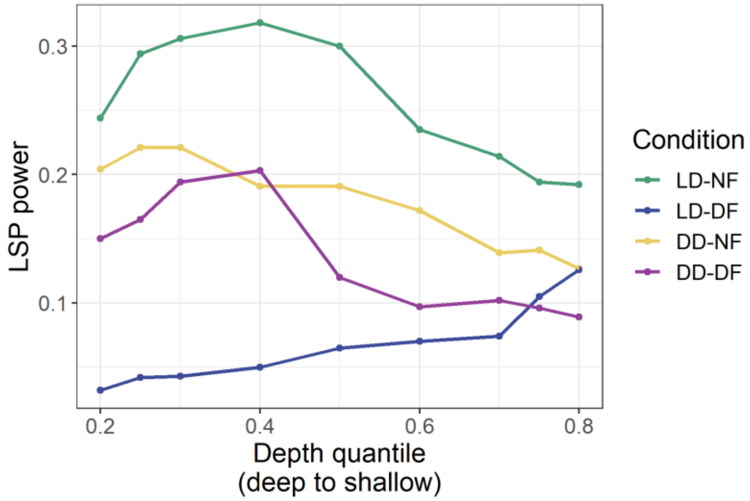


**Figure S2:** Periodogram power is higher in deeper depth quantiles. For each trial, depth quantiles were calculated, averaged into hourly bins, and LSP power computed (indicated by points in the figure). Lower quantiles occur deeper in the water column (e.g. the 0.25 quantile marks the deepest 25% of the group). Each point represents the Lomb-scargle periodogram power of a group's average time series for a given quantile.

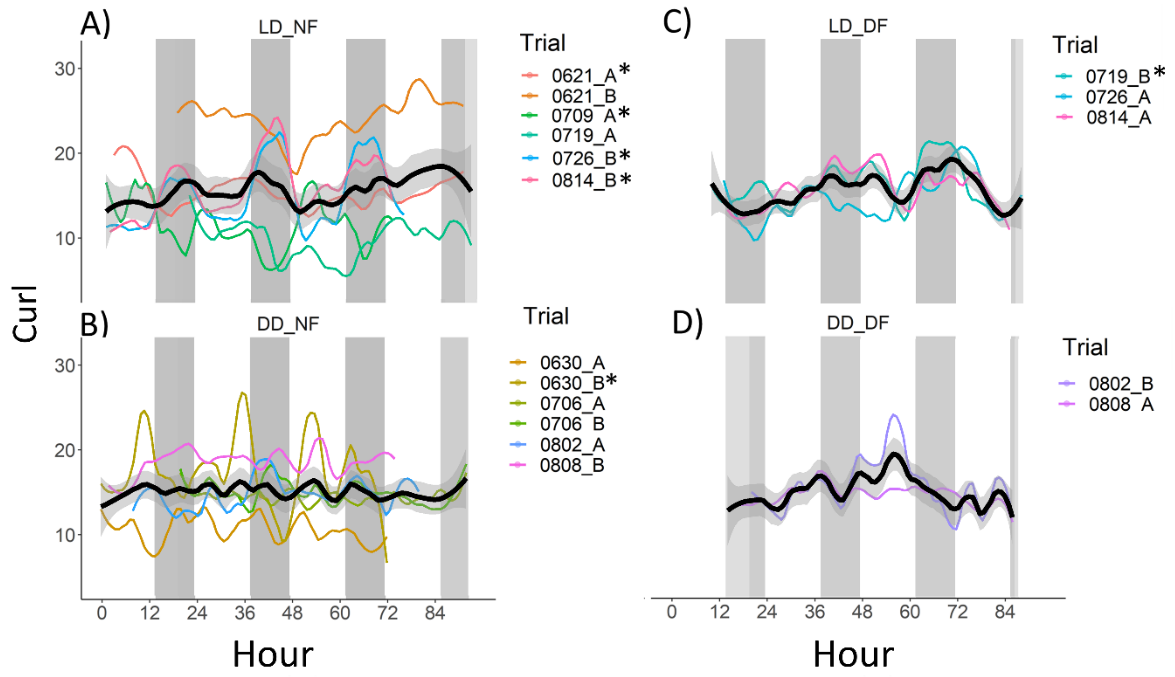

**Figure S3:** Curl over time. Colored lines represent loess smooths of hourly averages for each trial (points not shown for clarity). Black lines represent loess smooths of all data within each condition, and shaded areas represent 95% confidence intervals. A) LD cycle, nighttime feeding (LD-NF). B) Free-running, nighttime feeding (DD-NF). C) LD cycle, daytime feeding (LD-DF). D) Free-running, daytime feeding (DD-DF). Asterisks indicate trials with significant rhythmicity between 20–28h (LSP, p<0.01). Group average had significant diel rhythmicity only for LD-NF. White/grey rectangles represent lights-on/lights-off for panels A and B, and subjective day/night for panels C and D.

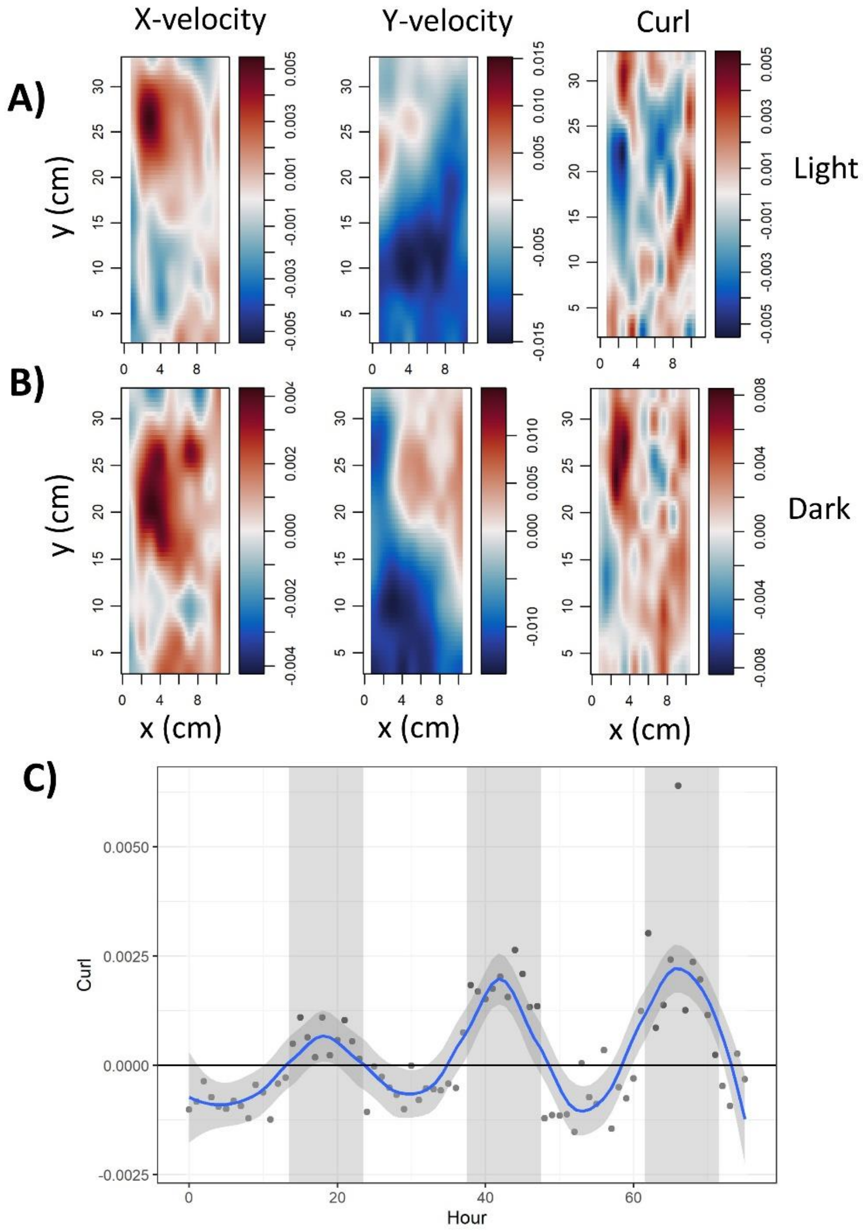


**Figure S4:** Example of trial where curl switches from positive to negative in a diel fashion (trial 0814-B). Trial was recorded in LD-NF conditions (Table 3A-1). A) X velocities (left), Y velocities (middle) and curl (right) averaged over lights-on hours. Positive values (red) indicate rightward movement, upward movement, and anti-clockwise rotation, respectively. B) X velocities (left), Y velocities (middle) and curl (right) averaged over lights-off (dark) hours. Positive values (red) indicate rightward movement, upward movement, and anti-clockwise rotation, respectively. C) Curl calculated within hourly bins and averaged over the entire tank. White/grey rectangles indicate lights-on/lights-off, the blue line is a loess smooth of the data, and the shaded area is the 95% confidence interval.

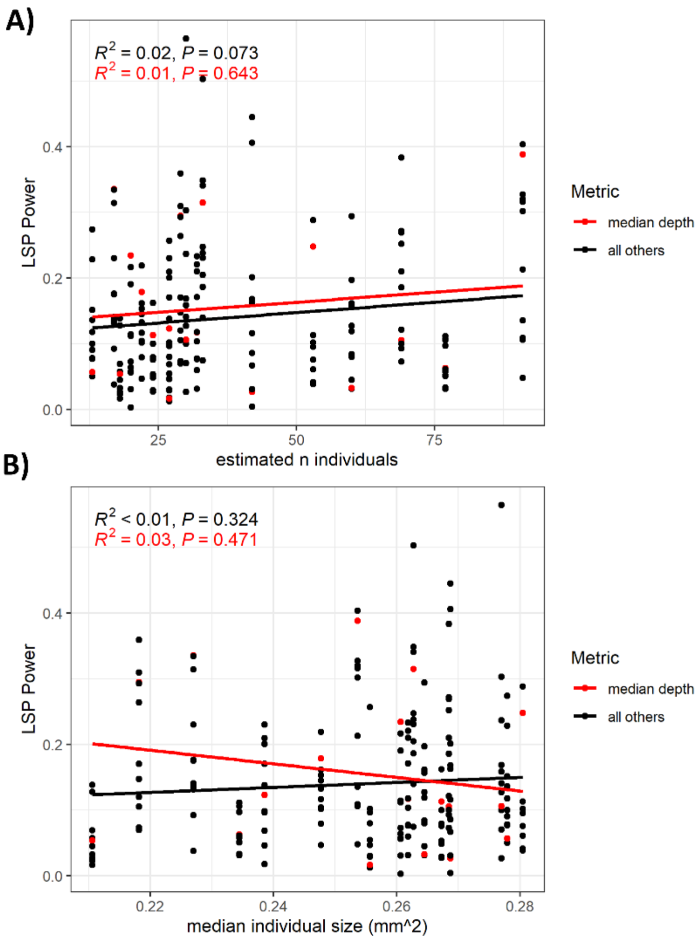


**Figure S5:** Strength of rhythmicity is not related to number of individuals (A) or median size (B). A) Number of individuals per trial was estimated as the maximum number of individuals tracked in at least 0.01% of frames (~50 frames). B) Median individual size per trial was calculated by considering all individual trajectories, regardless of length. Note that these sizes are only rough estimates and cannot be considered real measurements (see Methods). Each point represents periodogram power between 20-28 h for a particular trial and behavioral metric; see Table 1 for list of all metrics. Red points indicate power for median depth.
